# KIBRA-PKCγ signaling pathway modulates memory performance in mice and humans

**DOI:** 10.1101/2021.10.26.465926

**Authors:** Mengnan Tian, Qiang Chen, Austin R. Graves, Hana L. Goldschmidt, Richard C. Johnson, Daniel R. Weinberger, Richard L. Huganir

## Abstract

Human memory is a polygenic cognitive trait that is fundamental to individual competence. Genome-wide association studies (GWAS) have identified *KIBRA* as a novel gene associated with human memory performance. KIBRA interacts with AMPA receptors (AMPARs) and proteins essential for synaptic plasticity. The deletion of *Kibra* in mice impairs synaptic plasticity and learning and memory. However, the molecular basis through which KIBRA regulates dynamic AMPAR trafficking underlying synaptic plasticity is still unknown. Here we report that KIBRA interacts with the neuronal specific kinase PKCγ to modulate AMPAR trafficking upon learning, and KIBRA-PKCƔ signaling pathway also associates with human memory performance. We find PKCƔ is an essential kinase that phosphorylates AMPARs upon learning, and the loss of KIBRA in mouse brain impedes PKCƔ-AMPAR interaction. Activation of PKCƔ enables KIBRA to recruit phosphorylated AMPARs to the synapse to promote LTP and learning. We further performed transcriptomic and genetic analyses in human postmortem brain samples, and behavioral and fMRI evaluations in living human subjects, to demonstrate the genetic interactions between *KIBRA* and *PRKCG* on memory performance and memory associated physiological engagement of the hippocampal memory system. Overall, our results support that the KIBRA-PKCƔ signaling pathway is crucial for modulating memory performance in mice and humans.

---

Memory is critical for most higher brain functions and plays a central role in many behaviors across the phylogenetic spectrum. Decades of research have uncovered genes and signaling molecules that underlie learning and memory, many of which regulate synaptic plasticity^1,2^, the leading cellular mechanism for learning and memory. These essential genes and signaling molecules are often conserved across species^1^. Twin and familial studies in humans suggest that approximately half of the variability in memory performance can be attributed to genetic variation, especially in genes coding for synaptic proteins^2–4^. Until recently, memory-related molecules were implicated by linkage analysis or by candidate gene association studies^5^. However, these approaches were based primarily on pre-existing biological information, which biased investigation towards readily accessible molecular pathways and limited the potential of identifying novel memory-related genes. Genome-wide association studies (GWAS) avoid this potential bias by screening vast numbers of associations between heritable traits and millions of genetic variants distributed over the entire genome. This approach is particularly useful for discovering novel candidate genes and molecules that likely play critical roles in complex traits such as human cognition and disease. Furthermore, when these processes are conserved across species, this approach guides the development of animal models to investigate the molecular basis of cognitive disorders.

The first GWAS on human episodic memory identified the *KIBRA* gene as a potential regulator of human memory^6^, as carriers of a common C→T single nucleotide polymorphism (SNP rs17070145) within the ninth intron of *KIBRA* display improved episodic memory performance and hippocampal function^6^. Separate studies confirmed this association from subjects of different human populations across a wide age range^7–13^. KIBRA is a cytoplasmic scaffold protein highly enriched at the postsynaptic density^14^ (PSD), the crucial locus of synaptic communication between excitatory and inhibitory neurons. KIBRA interacts with memory-related molecules such as AMPA-type glutamate receptors (AMPARs), actin regulatory networks, and protein kinases^15,16^, but how these interactions contribute to cognitive performance is still unknown. In humans, the *KIBRA* T allele is also protective against Alzheimer’s disease (AD) risk^17,18^, and AD patients have decreased levels of KIBRA protein in the brain^19^. In mice, genetic deletion of *Kibra* impairs hippocampal synaptic plasticity and memory^14^, while over-expressing KIBRA rescues pathogenically disrupted synaptic plasticity in an AD mouse model^19^.

KIBRA may regulate memory by associating with proteins central to synaptic plasticity, such as AMPARs, which mediate the majority of fast-synaptic transmission in the mammalian brain^20^. Moreover, a recent study showed that membrane targeting of KIBRA is essential for long-term potentiation (LTP) and memory^21^. However, the molecular mechanism of how KIBRA regulates AMPAR membrane trafficking, synaptic plasticity, and memory remains completely unknown; and we are far from understanding the association between common *KIBRA* alleles and human cognitive performance. To address these fundamental issues, we examined the link between KIBRA expression and memory performance in mice and humans, investigating whether these mechanisms are conserved across species. Our data demonstrate that KIBRA engages PKCƔ to regulate AMPAR membrane trafficking, synaptic plasticity, and fear memory in mice. We also performed human genetic analysis and that the KIBRA and PKCƔ interaction is strongly associated with memory performance. Additionally, we confirmed that KIBRA recruitment to the neuronal membrane^21^ is associated with human memory performance as well. Our study elucidates the molecular interactions between KIBRA and several crucial synaptic proteins that orchestrate dynamic AMPAR trafficking underlying learning and memory, highlighting several novel potential therapeutic targets to enhance memory in normal humans and patients with cognitive disorders.

## Results

### KIBRA facilitates PKC interaction with AMPARs

To examine the molecular mechanism by which KIBRA regulates synapse function in the hippocampus, we used subcellular fractionation to study the localization of crucial synaptic proteins in WT and KIBRA knockout (KO) tissue. We isolated homogenate (Hom), postsynaptic density (PSD), and detergent extracted PSD (PSD II) fractions from mouse forebrain and probed for synaptic proteins. KIBRA is part of the AMPAR complex, but genetic deletion of KIBRA does not change the surface expression of the AMPAR GluA1/2 subunits^14^. However, KIBRA has been found to modulate protein kinase C (PKC) expression and function in various animals, ranging from Aplysia to mouse^22–26^. We found that KIBRA KO mice had decreased levels of multiple PKC isozymes, such as classical PKCα, β, γ, and atypical PKCι in the PSD II fraction (Fig 1A, B), suggesting that KIBRA is required for activation and the synaptic targeting of these PKC isozymes^27^. We also observed decreased PKMζ levels in the hippocampus Hom, PSD, and PSD II fractions in the KO mouse, consistent with previous results that PKMζ protein is not stable in the absence of KIBRA^26^. Next, we examined if decreased PKC membrane targeting inhibits PKC activity by probing PKC phosphorylation sites in the KIBRA KO hippocampus (Fig 1C, D). PKC phosphorylates the AMPAR GluA1 subunit at S831 (GluA1^S831^) and the GluA2 subunit at S880 (GluA2^S880^) residues, while PKA phosphorylates GluA1 at S845 (GluA1^S845^). We found that the phosphorylation of GluA1^S831^ and GluA2^S880^ were significantly lower in the KIBRA KO hippocampus lysate (GluA1^S831^: 83.61 ± 5.94% of WT littermate, *p* = 0.0185, n = 9; GluA2^S880^: 73.54 ± 4.72% of WT littermate, *p* = 0.0196, n = 9), but the phosphorylation of GluA1^S845^ was the same compared to that of WT littermates (103.66 ± 11.78% of WT, *p* = 0.7832, n = 9; Fig 1C, D). We also examined the phosphorylation of Tau on S321 (Tau^S231^), a PKC phosphorylation site on Tau^28^. We found that phosphorylation of Tau^S231^ was also reduced in the KIBRA KO hippocampus lysate (71.80 ± 9.34% of WT littermate, *p* = 0.0203, n = 9). Together, these data show that the loss of KIBRA significantly attenuates PKC signaling in the hippocampus.

**Fig. 1.**
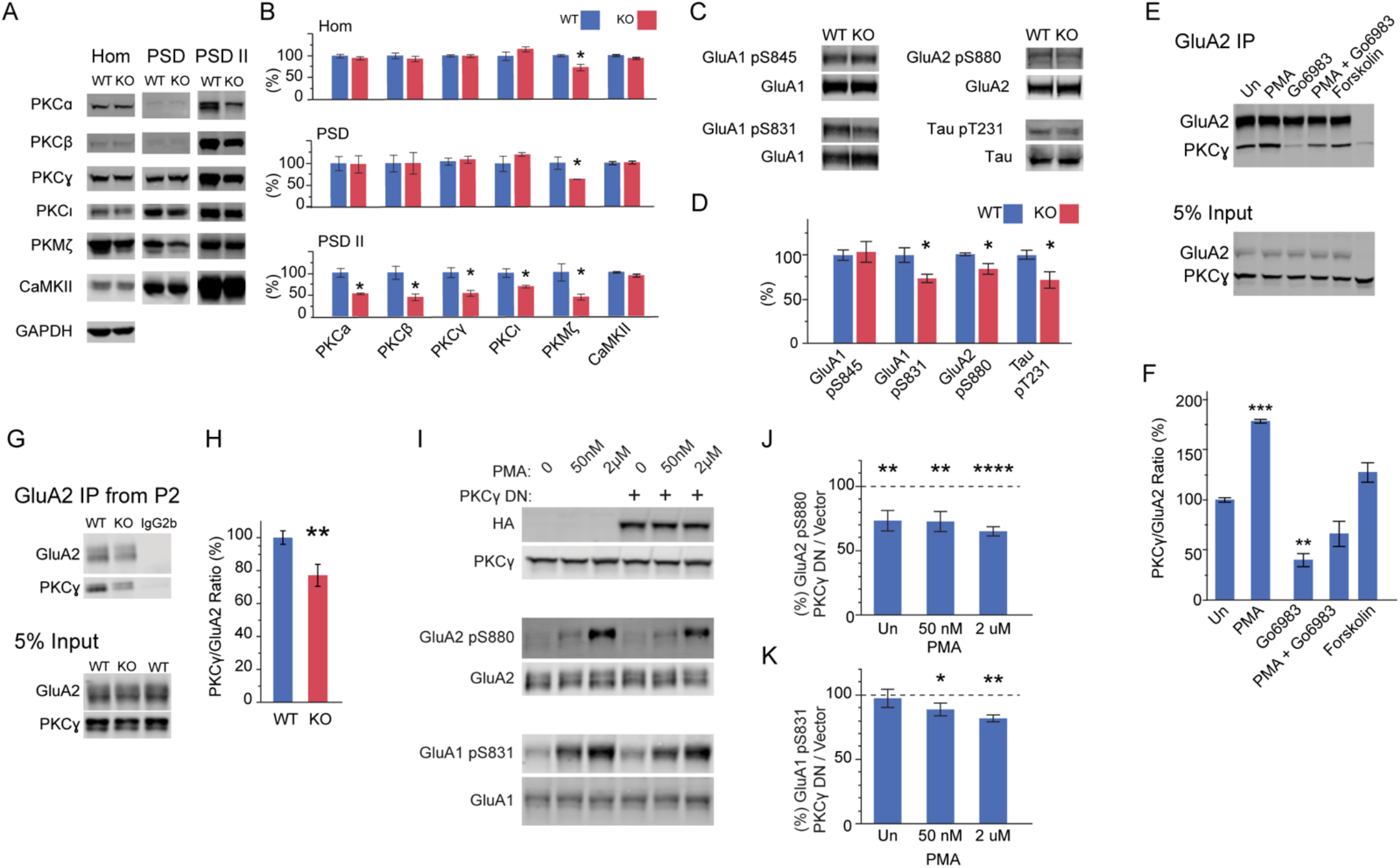
KIBRA facilitates PKCγ interaction with AMPARs. **A**. Biochemical fractionation of adult mouse forebrain reveals PKC family is decreased in KIBRA KO mouse forebrain postsynaptic density (PSD) fraction. Tissue homogenate (Hom), 0.5% Triton-X extracted postsynaptic density (PSD), and higher detergent extracted postsynaptic density (PSD-II) fractions were collected for Western blot. 10 mg of protein was loaded for each fraction. **B** Quantification of **A**. The value of each protein was compared to the mean of WT. n = 10, comparisons with control using Dunnett’s method. **C**. Western blot probing for phosphorylation level of protein kinase A and C sites in mouse hippocampus lysate. **D**. Quantification of **C**. n = 9, comparisons with control using Dunnett’s method. **E**. Representative western blots showing co-immunoprecipitation of PKCγ with GluA2 when co-expressed in HEK293T cells. Eighteen hours after transfection, HEK293T cells were treated with the chemical as labeled for 5 minutes and lysed in lysis buffer containing 1% NP40 and 0.5% Deoxycholate Sodium. PMA 2 μM, Go6983 10 μM, Forskolin 50 μM. **F**. Quantification of **E**. n = 4. One-way ANOVA *P* < 0.0001. **G** Representative western blots showing co-immunoprecipitation of PKCγ with GluA2 from P2 fraction of mouse forebrain. **H**. Quantification of **G**. n = 6. **I**. Overexpressing DN form of PKCγ inhibited AMPAR phosphorylation by PKC. Cultured neurons were electroporated with DN PKCγ and treated with 50 nM or 2 μM PMA for 5 minutes at DIV18. The phosphorylation level of GluA2 pS880 and GluA1 pS831 was probed and compared to that of vector electroporated neurons. **J** and **K** Quantification of **I**. Test of Means, n = 6. \**p* < 0.05; \*\**p* < 0.01; **** *p* < 0.0001.

PKC signaling is important for long-term potentiation (LTP), an essential mechanism underlying learning and memory^29^. While multiple PKC isozymes require KIBRA for membrane targeting and activation, not all are required for learning and memory^30–32^. At the molecular level, learning and memory are mediated by sustained changes in the number and activity of AMPARs in the postsynaptic membrane. To determine which PKC isozyme may be involved in KIBRA-regulated AMPAR trafficking, we first tested their ability to interact with the GluA2 AMPAR subunit and KIBRA. We co-expressed cDNAs encoding the GluA2 subunit and each PKC isozyme in HEK293T cells and performed co-immunoprecipitation with antibodies against the GluA2 subunit. Of the five isozymes tested, PKCγ was abundantly detected in GluA2 precipitates (Fig 1E, F). Importantly, the association of PKCγ with GluA2 was regulated by PKC activity (Fig 1E, F). The GluA2 subunit pulled down significantly more PKCγ in the presence of PKC agonist PMA and much less in the presence of a PKC inhibitor Go6983 (PMA: 178.34 ± 2.30% of untreated condition, *p* = 0.0002; Go6983: 37.66 ± 7.80% of untreated condition, *p* = 0.0015; n = 4, One-way ANOVA with Dunnett’s correction, *P* < 0.0001.). Treating cells with PMA in the presence of Go6983 did not affect GluA2/PKCγ interaction compared to untreated control (*p* = 0.1527, n = 4) or Go6983 treatment (*p* = 0.2601, n = 4), but much less compared to PMA treatment (*p* < 0.0001, n = 4). The adenylyl cyclase activator Forskolin had no effect on the GluA2/PKCγ interaction (*p* = 0.2730, n = 4). PKCγ also interacts with KIBRA (Supplement Fig 1A), but this interaction is not dependent on PKCγ activity. Further investigation showed both KIBRA and PKCγ bind to GluA2 subunit at the GluA2 C-terminus tail (Supplement Fig 1B, C). Deleting this region in the GluA2 subunit strongly decreased both GluA2/PKCγ and GluA2/KIBRA interactions.

To examine whether KIBRA is involved in the interaction between PKCγ and GluA2 subunit, we next examined PKCγ/KIBRA/GluA2 complexes in vivo. We collected P2 and PSD fractions from WT and KIBRA KO mouse forebrain, pulled down the GluA2 subunit, and determined the amount of PKCγ pulled down by GluA2 (Fig 1G, H, and Supplement 1D, E). Compared to WT littermates, the GluA2 subunit pulled down significantly less PKCγ from both P2 and PSD fractions in KIBRA KO mouse forebrain (P2: 77.10 ± 6.66% of WT littermate, *p* = 0.0099; PSD: 74.75 ± 7.94%, *p* = 0.0375; n = 6). To further investigate whether PKCγ directly phosphorylates AMPARs, we expressed a dominant-negative (DN) PKCγ in cultured neurons to inhibit PKCγ activity, treated the neurons with PKC agonist PMA, and then determined PKC phosphorylation at GluA2^S880^ and GluA1^S831^ residues by Western blot (Fig 1 I-K). Inhibition of PKCγ significantly impaired GluA2^S880^ and GluA1^S831^ phosphorylation, even at the saturating level of 2 μM PMA treatment. Consistent with the important role of PKCγ in modulating synaptic plasticity, learning, and memory^33–36^, these results combined indicate that PKCγ phosphorylates AMPARs and KIBRA can facilitate its interaction with AMPARs.

### KIBRA and active PKCγ are required for LTP

KIBRA KO mice have impaired hippocampus-dependent learning and memory upon trace auditory fear conditioning^14^. To understand the cellular mechanisms behind this cognitive deficit, we first used forskolin to induce a form of chemical LTP (cLTP) in cultured WT and KIBRA KO neurons to examine the role of KIBRA in recruiting AMPARs to the synapse upon LTP. We observed a 20%–30% increase in synaptic GluA1–3 subunits and AMPAR interacting protein GRIP1 in PSD fractions upon cLTP induction (Fig 2A-C). Interestingly, we also detected a significant increase in the abundance of KIBRA in the PSD (207.48 ± 39.03%, *p* = 0.0276, n = 7), substantially larger than the increase in AMPARs (Fig 2A-C). Notably, we saw no changes in total cellular levels of these proteins before and after cLTP, indicating selective enrichment of KIBRA, GRIP1, and AMPARs localized to the PSD. Remarkably, this PSD enrichment is abolished in KIBRA KO neurons (Fig 2A-C). During synaptic plasticity, increased intracellular Ca^2+^ levels result in CaMKII and PKC activation, directly phosphorylating different sites on AMPARs^20^. Consistent with this, we observed that levels of GluA1^S831^ phosphorylation increased to 134.40 ± 11.72% (*p* = 0.0301, n = 7) in Hom and 127.20 ± 7.38% (*p* = 0.0109, n = 7) in PSD fractions in cultured neurons following cLTP (Fig 2, D, E). cLTP induction also increased GluA1^S831^ phosphorylation in Hom fraction of cultured KIBRA KO neurons to 126.79 ± 8.88% of WT neurons but not in the PSD fraction (Hom: p = 0.0322; PSD: *p* = 0.3439; n = 7) (Fig 2F, G). These data demonstrate that KIBRA plays a critical role in the recruitment of AMPAR subunits GluA1-3, GRIP and GluA1^S831^ to the PSD during chemical LTP.

**Fig 2.**
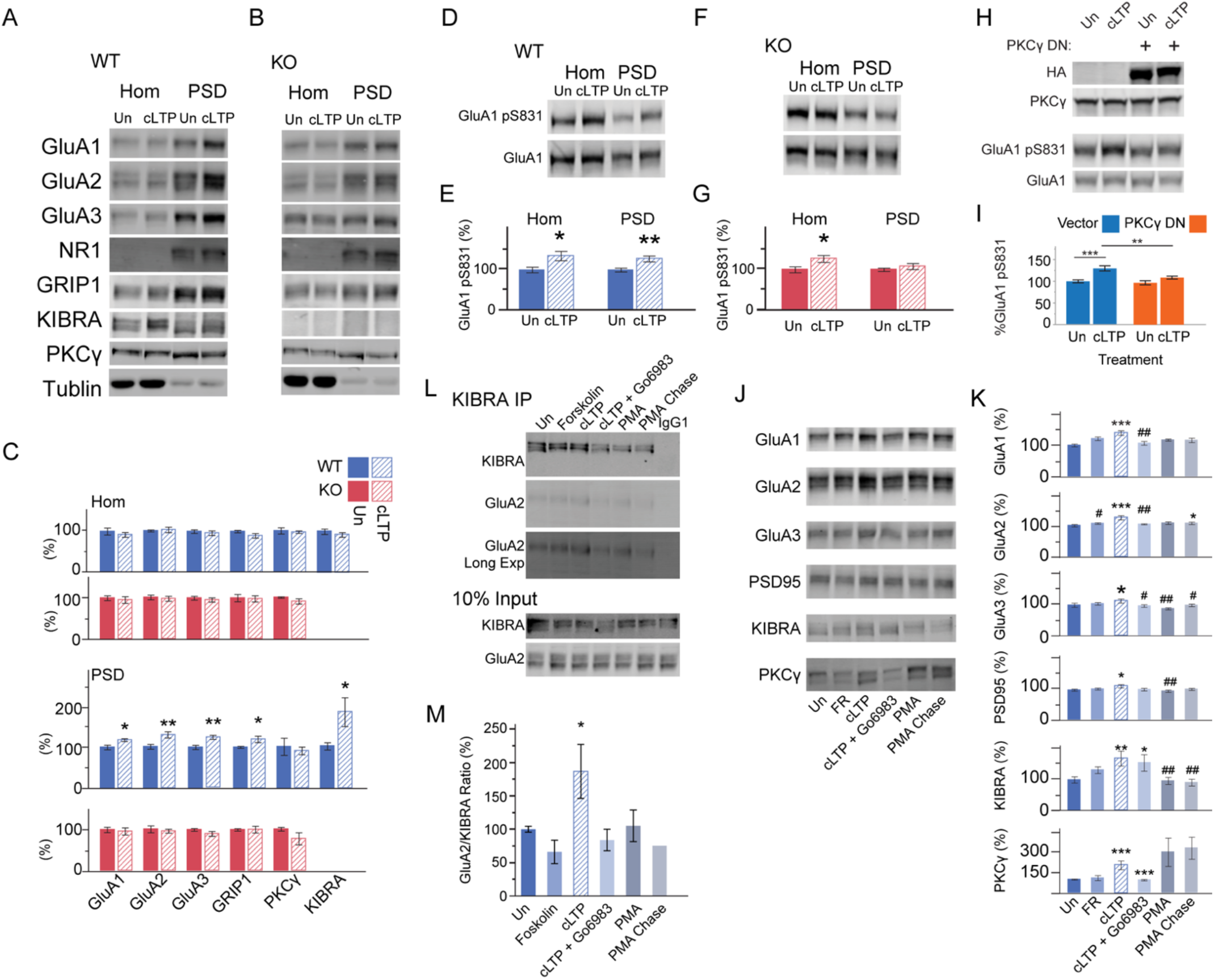
KIBRA, PKCγ, and AMPAR interaction enhanced upon induction of synaptic plasticity in cultured mouse hippocampal neurons. **A** and **B**. Representative western blots of proteins from Hom and PSD isolated from cultured WT and KIBRA KO neurons before and after forskolin cLTP induction. **C** Quantification of **A** and **B**. WT: n = 7; KO: n = 11. **D**. Representative western blots of proteins from Hom and PSD isolated from cultured WT neurons before and after forskolin cLTP induction. **E** Quantification of **D**. n = 7. **F**. Representative western blots of proteins from Hom and PSD isolated from cultured KIBRA KO neurons before and after forskolin cLTP induction. **G.** Quantification of **F**. n = 7. **H**. Representative western blots of Hom and PSD proteins isolated from culture neuron expressing vector or DN PKCγ at DIV19. **I**. Quantification of **H**. n = 10. **J**. Representative western blots of PSD proteins isolated from cultured neurons treated with chemicals. Forskolin: 50 μM forskolin with 0.1 μM rolipram treat for 15 minutes; cLTP: Forskolin treatment followed by 45 minutes chase; cLTP+Go6983: cLTP in the presence of 10 μM Go6983; PMA: 2 μM treat for 15 minutes; PMA Chase: PMA treatment followed by 45 minutes chase. **K.** Quantification of **J**. n = 8. *, # *p* < 0.05; **, ## *p* < 0.1; \*\*\**p* < 0.001. # compared to cLTP. **L**. Representative western blots co-immunoprecipitation from cultured neurons treated with chemicals. Forskolin: 50 μM forskolin with 0.1 μM rolipram treat for 15 minutes; cLTP: Forskolin treatment followed by 45 minutes chase; cLTP + Go6983: cLTP in the presence of 10 μM Go6983; PMA: 2 μM treat for 15 minutes; PMA Chase: PMA treatment followed by 45 minutes chase. **M.** Quantification of **L**. n = 10.

Next, we explored the role of PKCγ in AMPAR phosphorylation during LTP by inhibiting PKCγ in cultured neurons. Upon cLTP stimulation, GluA1^S831^ phosphorylation increased 129.68 ± 5.88% (*p* = 0.0004, n = 10) in vector-expressed neurons but not in neurons expressing dominant-negative (DN) PKCγ (*p* = 0.0585, n = 10) (Fig 2H, I). These data demonstrate that PKCγ is a major PKC isozyme that phosphorylates AMPARs upon cLTP. To further understand the role of PKC activity in LTP, we pharmacologically manipulated PKC activity during cLTP in cultured neurons. AMPAR subunits GluA1-3 were enriched about 30% in the PSD following cLTP, but this increase was not observed in the presence of the PKC inhibitor Go6983 (Fig 2J, K). Surprisingly, although KIBRA and PKCγ were also enriched upon cLTP, Go6983 blocked PKCγ enrichment upon cLTP but did not affect KIBRA enrichment (Fig 2J, K). Treating neurons with PMA increased the PKCγ level at PSD but did not increase KIBRA or AMPAR levels. KIBRA binds to AMPARs at GluA2 C-terminus tail (Supplement Fig 1B), so we also pulled down KIBRA from neuron lysates and probed for the GluA2 subunit to examine KIBRA/AMPAR interaction (Fig 2L, M). KIBRA pulled down twice as much GluA2 upon cLTP (*p* = 0.0009, n = 10, One-Way ANOVA with Dunnett’s correction, *P* = 0.0004), and this increase was blocked in the presence of Go6983. PMA treatment activated PKCs but did not increase KIBRA/GluA2 interaction. Taken together these results suggest that both KIBRA translocation to the PSD and PKC activation are required for LTP, where KIBRA translocation to synapses upon LTP recruits PKCγ-phosphorylated AMPARs to synapses.

### KIBRA and PKCγ are enriched at the PSD following learning

We next tested whether PKCγ regulates KIBRA-dependent AMPAR trafficking during mouse learning and memory. Using hippocampus-dependent contextual fear conditioning in KIBRA KO mice (Fig 3A, B), we found that KIBRA KO mice showed significantly less freezing than WT littermates 24 hours (*p* = 0.0068, n = 8) and one week (*p* = 0.0344, n = 8) after training (Fig 3C). We collected the hippocampus from WT mice 2 hours after training and probed for synaptic protein expression in Hom and PSD fractions. We observed a 20% increase in synaptic GluA1–3 subunits in WT mice PSD fractions upon fear learning compared to home cage control (Fig 3D, E). Consistent with previous reports, we found that fear conditioning increased CaMKIIɑ in the PSD fraction (161.71 ± 11.74%, *p* = 0.0008, n = 9). Autophosphorylation of CaMKIIɑ^T286^ was similarly increased (166.62 ± 21.23% in Hom fraction, *p* = 0.0192, n = 9; 117.18 ± 3.88% in PSD fraction, *p* = 0.1134, n = 9). Interestingly, we detected a significant increase in the abundance of KIBRA and PKCγ in the PSD (KIBRA: 139.19 ± 13.40%, *p* = 0.0304; PKCγ: 148.65 ± 15.16%, *p* = 0.0088; n = 9). Consistent with cLTP results, we saw no change in total cellular levels of these proteins before and after fear learning, suggesting that PSD enrichment of KIBRA, PKCγ, AMPARs, and CaMKIIɑ are a result of specific synaptic trafficking events rather than an effect on total protein levels (Fig 3D, E). Next, we examined synaptic protein expression in KIBRA KO mice and WT littermates before and after fear learning. KIBRA KO mice had normal levels of AMPARs, CaMKIIɑ, and PKCγ in hippocampus Hom and PSD fractions before fear learning (Supplementary Fig 1F, G). However, levels of GluA1-3 in KIBRA KO hippocampus PSD were significantly lower than those of WT littermate 2 hours after fear learning (Fig 3F, G). This result indicates that AMPARs enrichment in PSD after learning (Fig 3D, E) requires KIBRA, as observed in cultured KO neurons upon cLTP (Fig 2A-C). Notably, PKCγ in KO PSD after fear learning was also significantly lower than WT littermates (65.71 ± 8.04%, *p* = 0.0027; n = 9), suggesting that PSD translocation of PKCγ depends on KIBRA.

**Fig 3.**
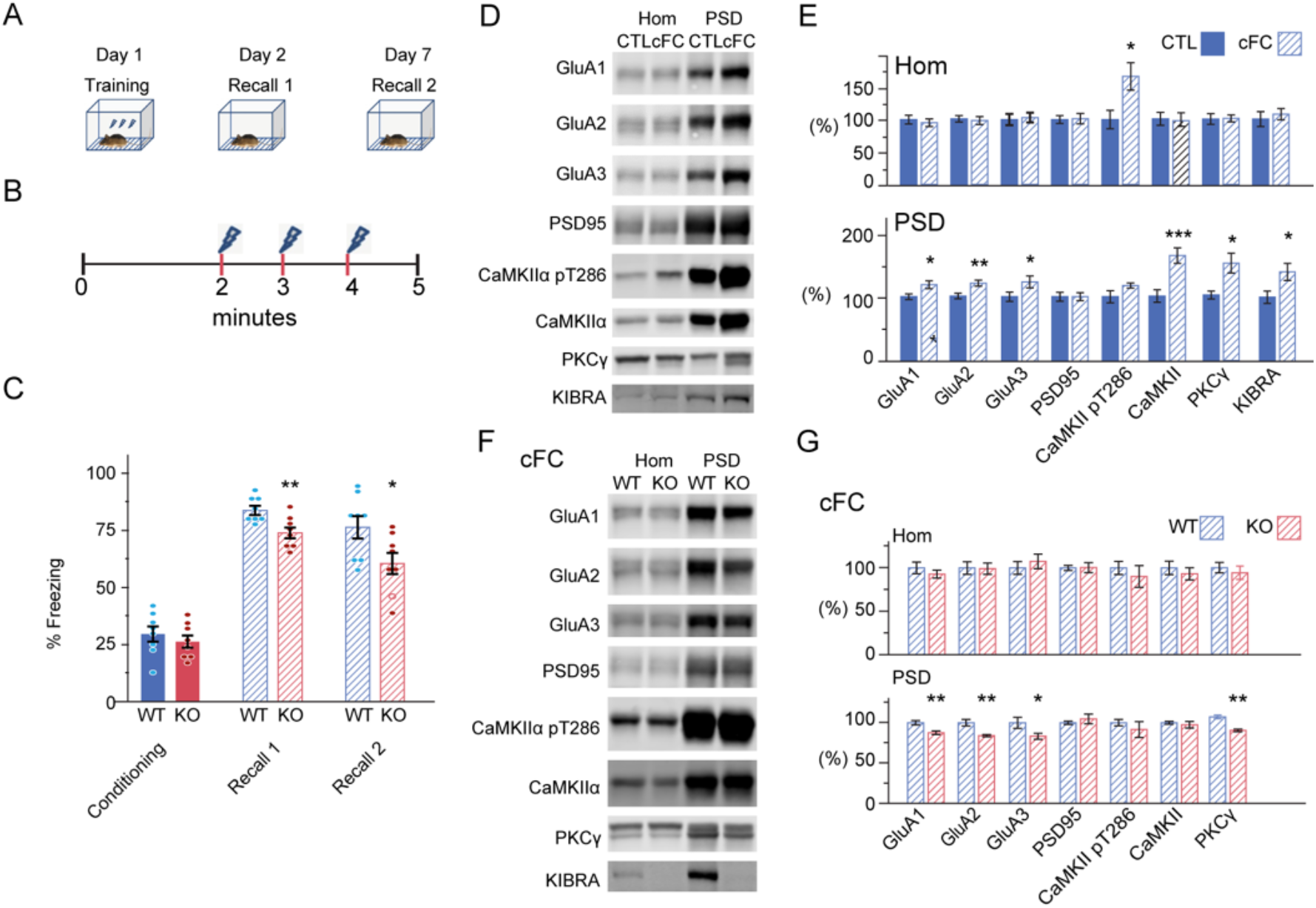
KIBRA is crucial for AMPAR complex and PKCγ enrichment on PSD upon mouse fear learning. **A** and **B**. Experiment design for behavior training. **C**. Analysis of fear-related freezing showing KIBRA KO mice had less freezing during two recall sessions. n = 8 mice. **D**. Representative western blots of tissue homogenate (Hom) and postsynaptic density (PSD) proteins from the hippocampus of WT mice with or without fear learning. **E**. Quantifications of **D**. n = 9. CTL: Home cage control; cFC: contextual fear learning. **F**. Representative western blots of tissue Hom and PSD proteins from the hippocampus of gender-matched WT and KIBRA KO littermates after fear learning. **G**. Quantifications of **F**. After fear learning, KIBRA KO mice had significantly fewer AMPARs and PKCγ than WT littermate. n = 10. \**p* < 0.05; \*\**p* < 0.01; \*\*\**p* < 0.001

However, fear learning still activated CaMKIIɑ in the KIBRA KO mouse hippocampus, as the increase of CaMKIIɑ and CaMKIIɑ^T286^ phosphorylation in the PSD was the same as WT. Together, these results suggest that KIBRA is not required for CaMKIIɑ translocation and phosphorylation but is essential for AMPAR recruitment to the PSD during learning.

Consistent with cLTP, we observed that levels of GluA1^S831^ phosphorylation increased about 30-40% in both Hom and PSD fractions in mouse hippocampus after fear learning (Fig 4A, B). Home-cage control KIBRA KO mice had 20-25% lower of GluA1^S831^ phosphorylation in both Hom and PSD fractions compared to WT littermates (Hom: 78.91 ± 4.43%, *p* = 0.0024; PSD: 75.35 ± 5.21%, *p* = 0.0004; n = 10) (Fig 4C, D). After fear learning, GluA1^S831^ phosphorylation in KIBRA KO hippocampus Hom fraction was not different from that of the WT hippocampus, but still significantly lower in the KIBRA KO PSD fraction (Hom: 99.44 ± 7.39%, *p* = 0.9606; PSD: 76.27 ± 2.60%, *p* = 0.0011; n = 6, Dunnett’s comparison; Fig 4E, F). KIBRA KO mice had normal CaMKII activation upon fear learning, but PKCγ level in KIBRA KO PSD fraction after fear learning is also significantly less compared to WT PSD (Hom: 99.52 ± 4.93%, *p* = 0.9656; PSD: 65.71 ± 8.04%, *p* = 0.0027; n = 6, Dunnett’s comparison). We then pulled down GluA2 from WT forebrain before and after contextual fear learning and found it pulled down 129.38 ± 10.47% of PKCγ after fear learning compared to the control (*p* = 0.0178, n = 8) (Fig 4G, H). We also pulled down KIBRA from these tissues and found the amount of GluA2 and PKCγ co-immunoprecipitated with KIBRA increased to 143.82 ± 14.62% and 144.71 ± 10.89%, respectively, after learning (*p* = 0.0258 and 0.0244, n = 8) (Fig 4I-K). Taken together with cLTP results, these data demonstrated that PKCγ is a major PKC isozyme that phosphorylates AMPARs upon LTP and mouse learning. Its interaction with KIBRA and AMPARs is enhanced after learning, which increases phosphorylated AMPARs trafficking to the synapses in the presence of KIBRA.

**Fig 4.**
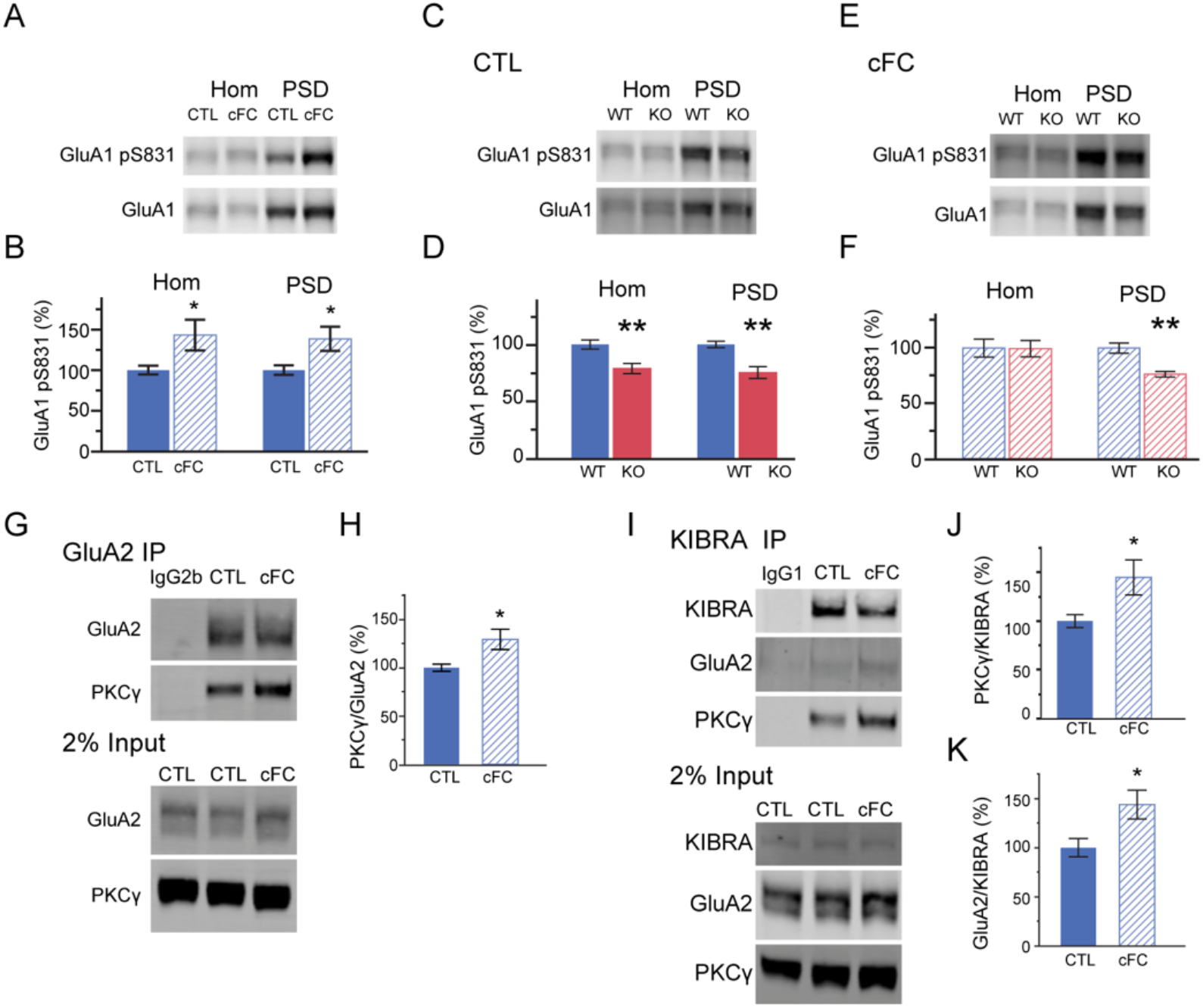
PKCγ is essential for KIBRA regulated AMPAR trafficking upon synaptic plasticity. **A**. Representative western blots of Hom and PSD proteins isolated from WT mouse hippocampus before and after contextual fear learning. **B.** Quantification of **A**. n = 8. **C**. Representative western blots of proteins from Hom and PSD isolated from the hippocampus of WT and KIBRA KO littermates in the home cage. **D.** Quantification of **C**. n = 10. **E**. Representative western blots of Hom and PSD proteins isolated from the hippocampus of WT and KIBRA KO littermates after contextual fear learning. **E.** Quantification of **F**. n = 6. **G**. Representative western blots showing co-immunoprecipitation of PKCγ with GluA2 from P2 fraction of WT mouse forebrain before and after contextual fear learning. **H**. Quantification of **G**. n = 8. **I**. Representative western blots showing co-immunoprecipitation of PKCγ and GluA2 with KIBRA from P2 fraction of WT mouse forebrain before and after contextual fear learning. **J.** and **K**. Quantification of **I**. n = 8. \**p* < 0.05; \*\**p* < 0.01.

### KIBRA is positively associated with human memory performance

KIBRA has been associated with brain function in mice, and *KIBRA* rs17070145 SNP associates with cognitive performances in humans^6–9,37^. Therefore, we tested whether the KIBRA and PKCγ signaling pathways may interact to influence human memory performance as well. Using neuropsychological test data for 308 participants with European ancestry (age: 18-86, 32.5 ± 12.82, mean ± SD), we first confirmed the association of *KIBRA* genotype with human memory test performance and with hippocampal activity determined with fMRI. Consistent with previous publications, the T-allele carriers have relatively better cognitive performance and relatively greater hippocampal engagement during an episodic memory task. (Supplement Fig 2A-F)^9^. We studied age-related human cognitive behavior and memory-related hippocampal activity differences as indicators of human memory performance, as previous studies suggested that the genetic impact of KIBRA is potentiated at older ages^9^.

To better understand the human neuropsychological and neuroimaging genetic association data, we mapped expression quantitative trait loci (eQTLs) in 395 postmortem human bulk hippocampus samples from the “Brain-seq” RNA sequencing dataset^38^ and found alleles of *KIBRA* rs17070145 are associated with varying levels of KIBRA mRNA expression in the human hippocampus (Fig 5A). Earlier studies grouped the heterozygous TC allele genotype individuals with the homozygous minor TT genotype individuals to balance the genotype contrast groups and compared the behavioral evaluations between the T-allele and CC-allele groups. Our eQTL analysis showed that KIBRA T-allele carriers had significantly greater mRNA expression than the CC-genotype group (*p* = 0.00107, Fig 5A). Thus, the *KIBRA* rs17070145 T-allele is associated with enhanced human memory performance, with greater hippocampal physiological engagement during episodic memory (Fig Supplement 2A-F) and also greater KIBRA mRNA expression in the human hippocampus (Fig 5A). Next, we found that SNP rs12151321, an eQTL of *PRKCG*, showed allelic association with the PKCγ mRNA level in the human hippocampus RNA seq data (Fig 5B) where A-allele carriers (grouped as AA-genotype plus AC-genotype individuals) have significantly lower PKCγ mRNA expression compared to CC-genotype carriers (*p* = 0.000767, Fig 5B). The A-allele carriers and CC-genotype samples of *PRKCG* rs12151321 did not show any differences in age-associated delayed recall performance or episodic memory-related hippocampus activity during fMRI (Supplementary Table 1, 2).

**Fig 5.**
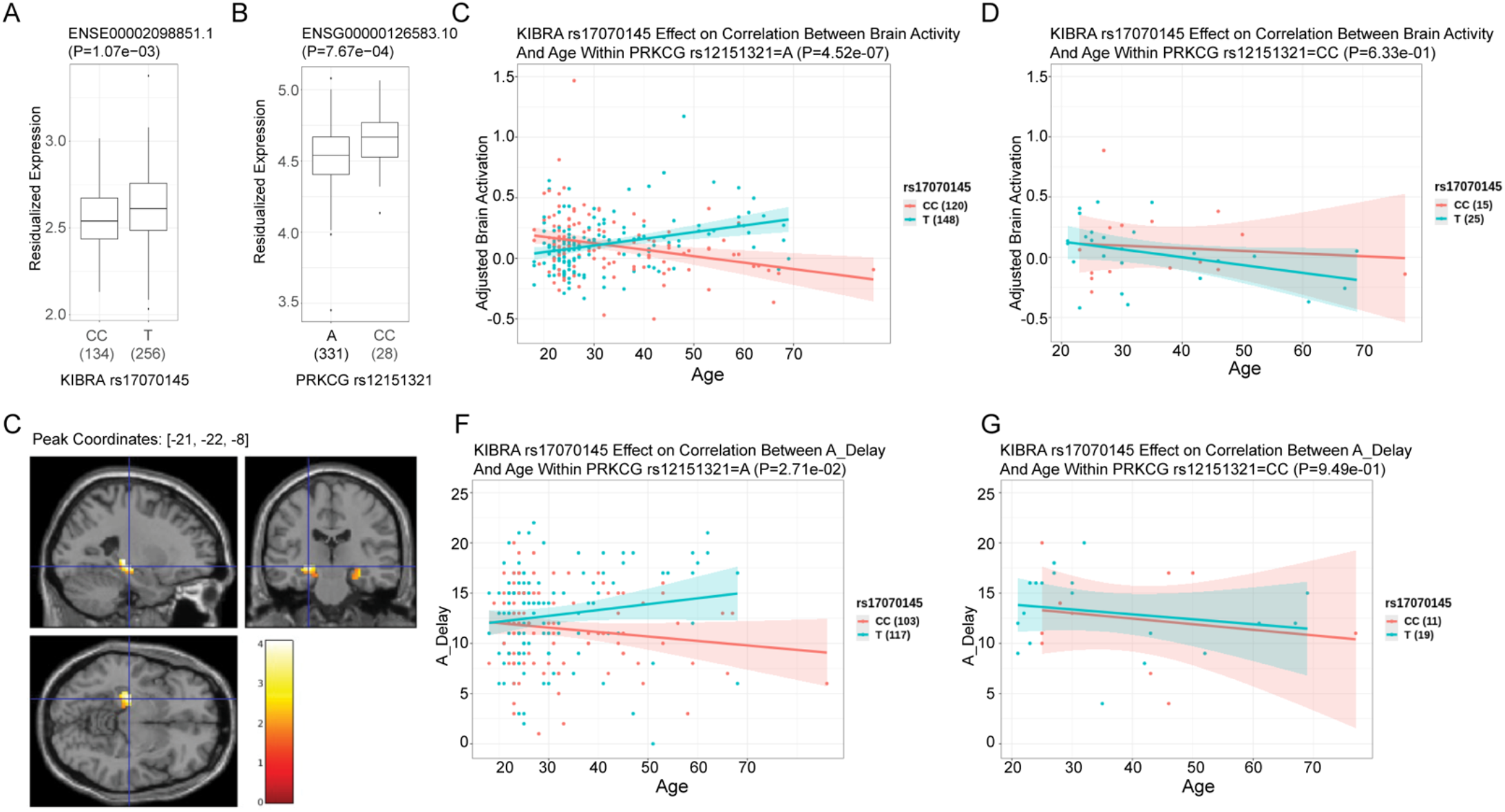
The allelic association of the *KIBRA* gene with human memory performance and brain activation under episodic memory task are modified by eQTLs of *PRKCG*. **A** and **B** eQTL effect on expression of *KIBRA* and *PRKCG*. **C-E:** Association of age with brain activation under episodic memory task affected by *KIBRA* SNP rs17070145 stratified by *PRKCG* SNP rs12151321. The difference between rs17070145 x age interaction effects in two rs12151321 genotypes is significant with P=0.0043. **C:** Location of interaction effect. **D, E:** Brain activation vs age in each genotype group. A_Delay: Delayed recall. **F, G:** Association of age with memory performance affected by *KIBRA* SNP rs17070145 stratified by *PRKCG* SNP rs12151321. The difference between rs17070145 x age interaction effects in two rs12151321 genotypes show a trend with P=0.1993.

We then explored the possibility that KIBRA and *PRKCG* would show interactive effects in the human data beyond their individual associations. We first examined whether variation in PKCγ mRNA expression levels can modulate the allele-dependent association between KIBRA and human memory performance. Remarkably, we found *PRKCG* rs12151321 and *KIBRA* rs17070145 genotypes showed strong interaction (P=0.0042) in human hippocampus activation during the episodic memory test (Fig 5C-E). The differential association of *KIBRA* rs17070145 alleles with age-related hippocampus activity during episodic memory was still observed within the *PRKCG* rs12151321 A group but not in the CC group (Fig 5C, D, Supplementary Table 2). Interestingly, we also found that although the differential association of *KIBRA* rs17070145 alleles with age-related delayed recall performance was still observed within the *PRKCG* rs12151321 A group, this association was not observed in the *PRKCG* rs12151321 CC group, although test for two SNPs interaction effect on age-related delayed recall performance is not significant (P=0.1993) (Fig 5F, G, Supplementary Table 1). Increased PKCγ mRNA expression in the *PRKCG* rs12151321 CC-carriers was associated with apparent rescue of the allelic differential association of *KIBRA* rs17070145 SNP with human memory performance. This suggests that the *KIBRA* rs17070145 CC-allele carriers association with worse memory performance, possibly due to lower KIBRA expression in the hippocampus, is compensated for by individuals with higher hippocampal PKCγ expression. In contrast, the eQTL at the *PRKCZ* gene did not modify *KIBRA* rs17070145 SNP association with human memory (Supplementary Fig 2G, Supplementary Table 1, 2), despite that KIBRA binds to PKMζ and stabilize it in the mouse brain (Fig 1A and B)^24–26,39^. Together with the in vitro and in vivo biochemical data, these data strongly support a model whereby PKCγ, in facilitating KIBRA-regulated AMPAR trafficking in the hippocampus during learning, underwrites memory function.

These results reveal a novel, phylogenetically conserved mechanism of the interaction of KIBRA with PKCγ to direct AMPAR trafficking and lead to synaptic plasticity and learning and memory. We first established that PKCγ regulated KIBRA-dependent AMPAR trafficking during LTP. Both fear learning and LTP stimulation activate PKCγ, which phosphorylates AMPARs and increases AMPAR expression in the PSD via KIBRA/GluA2 interaction. We then performed human eQTL analysis and examined the neuropsychological performance and episodic memory-related hippocampus activity. The results showed that while *PRKCG* gene does not associate with human memory performance alone, increased PKCγ in the human hippocampus may compensate for the lower KIBRA mRNA expression and abolished the allele dependent KIBRA association with human memory performance. These results indicate KIBRA is a crucial protein modulating memory performance, and its function depends on interactions with PKCγ. Further, these novel mechanistic insights highlight many new targets for clinical interventions and pharmacological treatments of a wide array of neuropsychiatric diseases.

## Methods (Including Separate Data And Code Availability Statement)

### Subjects

All animals were treated following the Johns Hopkins University Animal Care and Use Committee guidelines. Wild-type and KIBRA KO mice were of the C57BL6 hybrid background. Gender-matched WT and KIBRA KO littermates were randomly assigned to experimental groups and used at embryonic day 18 (E18), juvenile (3 weeks), or adult (3–6 months) stage as indicated. Sprague Dawley rats were used for E18 neuron cultures. All animals were group-housed in a standard 12-hour light/ 12-hour dark cycle.

HEK293T cells were transfected using Lipofectamine2000 (Invitrogen) according to manufacture recommendation. Rat neurons were transfected using Rat Neuron Nucleofector® Kit (Lonza Group) according to manufacture recommendation.

### Contextual fear conditioning

Mice were handled for 5 min on each of 5 consecutive days before beginning experiments. The training chamber was located inside a sound-attenuating isolation box and consisted of a square chamber with two Plexiglas and 2 aluminum sides, and a remote-controlled electrifiable grid floor. Training occurred on Day 1 as follows: Mice were allowed acclimate to the chamber for 2 min before the onset of training blocks, then received three electric foot shocks (0.75 mA, 2 s duration) at 1-minute intervals. Mice were returned to their home cage immediately following training. On Day 2 or Day 7, mice were placed in the training chamber for 5 min to assess contextual fear conditioning, after which they were returned to their home cage. The percentage of time freezing was quantified using automated motion detection software (AnyMaze, Stoelting Co). For Figure 2C, a two-way ANOVA (genotype x interval) was used to detect significant differences between groups, and significant differences at specific intervals were established with a Tukey post-hoc test. In all other cases, a two-tailed student’s t-test was used.

### Forskolin-induced LTP and drug treatment

For cLTP experiments, cultured cortical neurons at DIV17-19 were treated with 50 μM Forskolin, 0.1 μM rolipram in culture media for 15 minutes, and then returned to the culture media for 45 minutes chase before lysis or homogenate for co-immunoprecipitation or PSD preparation. For Go6983 inhibition, 10 μM Go6983 was co-applied with Forskolin/rolipram and added to culture media during 45 minutes chase. PMA was applied at 2 μM for 15 minutes, or 50 nM and 2 μM for 5 minutes in Figure 1I-K.

### PSD subcellular fractionation

Cultured cortical neurons or mouse hippocampus tissue was homogenized by passage through a 26g needle, 12 times, in homogenization buffer (320mM sucrose, 5mM sodium pyrophosphate, 1 mM EDTA, 10 mM HEPES (pH 7.4), 200 nM okadaic acid, and protease inhibitors). The homogenate was centrifuged at 800xg for 10 minutes at 4°C to yield post-nuclear pelleted fraction <u>1</u> (P1) and supernatant fraction <u>1</u> (S1). S1 was further centrifuged at 15,000xg for 20 minutes at 4°C to yield P2 and S2. P2 was resuspended in milliQ water, adjusted to 4 mM HEPES (pH 7.4) from a 1 M HEPES stock solution, and incubated with agitation at 4°C for 30 minutes. The suspended P2 was centrifuged at 25,000xg for 20 minutes at 4°C to yield LP1 and LS2. LP1 was resuspended in 50 mM HEPES (pH 7.4), mixed with an equal volume of 1% Triton X-100, and incubated with agitation at 4°C for 15 minutes. The PSD was generated by centrifugation at 32,000xg for 20 minutes at 4°C. The final PSD pellet was resuspended in RIPA buffer (1% NP-40, 0.5% Sodium Deoxycholate, 0.1% SDS, 200 mM NaCl, 50 mM sodium fluoride, 5 mM sodium pyrophosphate, 20 mM HEPES (pH 7.4), and protease inhibitors) followed by protein quantification and Western blot.

Mouse forebrain tissue was homogenized with a Teflon glass homogenizer for 35 strokes, and then the sample was centrifuged 10 min at 1,000 × *g*, and the supernatant was centrifuged at top speed for 20 min. The resulting pellet (P2) was resuspended in homogenization buffer and layered onto a discontinuous sucrose gradient containing 0.85 M/1.0 M/1.4 M sucrose, and centrifuged at 82,500 × g for 2 h. The band at the 1.0 M/1.4 M sucrose interface was collected and centrifuged at 150,000 × g for 30 min. The resulting pellet (synaptosome; SPM) was solubilized with 0.5% Triton X-100 solution (containing 40 mM Tris·HCl pH = 8.0 and protease inhibitor mixture) and centrifuged at 32,000 × g for 20 min to yield PSD pellet.

### Coimmunoprecipitations and Immunoblotting

HEK cells, cultured neurons, or mouse brain fractions were lysed in lysis buffer (1% NP-40, 0.5% Sodium Deoxycholate, 200 mM NaCl, 50 mM sodium fluoride, 5 mM sodium pyrophosphate, 20 mM HEPES (pH 7.4), and protease inhibitors) for 20-60 minutes at 4°C with agitation, then the lysates were collected and centrifuged at maximum speed in the cold room for 15 minutes. 0.5-1 mg protein was first incubated with 5-10 μg purified antibodies against KIBRA or GluA2 at 4°C overnight, followed by adding 20 μl Protein G beads for 2 hours. Beads were washed four times with lysis buffer without protease inhibitors, and immune complexes were eluted in a 2x NuPage sample buffer 10 min. Eluates were resolved by 8 or 10% SDS-polyacrylamide gel electrophoresis (PAGE), transferred to PVDF membrane, and immunoblotted to examine protein of interest using a specific primary antibody. The corresponding secondary antibodies used were IRDye-coupled whole antibodies (LI-COR Biosciences). The Odyssey Infrared Imaging System protocols were used for blots and quantification of protein expression.

### Quantification and statistical analysis

All data were analyzed in JMP (SAS Institute) and presented as mean ± SEM (standard error of the mean) unless indicated otherwise. All statistical details and statistical significance calculated using the Mann-Whitney test, Kolmogorov-Smirnov test, unpaired t-test, one-way or two-way ANOVA, were indicated the figure legends. Tukey and Bonferroni post-hoc tests were used following one-way and two-way ANOVA, respectively. *, p<0.05; **, p<0.01, ***, p<0.001.

### Human Postmortem Brain Sample

Postmortem human brain tissue was obtained by autopsy primarily from the Offices of the Chief Medical Examiner of the District of Columbia and of the Commonwealth of Virginia, Northern District, all with informed consent from the legal next of kin (protocol 90-M-0142 approved by the NIMH/NIH Institutional Review Board). Details of tissue acquisition, handling, processing, dissection, clinical characterization, diagnoses, neuropathological examinations, RNA extraction and quality control measures were described previously by Lipska et al^40^. The Brain and Tissue Bank cases were handled in a similar fashion (http://www.medschool.umaryland.edu/btbank/Medical-Examiners-and-Pathologists/Minimum-Protocol/). Hippocampus sample processing was described by Collado-Torreet et al^38^.

### Human Postmortem eQTL Analysis

Details of RNA data processing, genotype data processing, and eQTL analysis were previously described by Collado-Torreet et al^2^. The related gene expression and eQTL results for hippocampus could be found at BrainSeq Consortium Website: http://eqtl.brainseq.org/

### Human Postmortem SNP Selection

Based on eQTL results in hippocampus from the BrainSeq Consortium, we used the clumping approach in PLINK (version 1.9, https://www.cog-genomics.org/plink/) to remove SNPs in LD and obain the top index eQTLs with FDR<0.05 in each clump that includes at least 3 SNPs. These top index eQTLs were used for further analysis using human clinical samples. SNPs were converted to two goups according to expression levels in three genotype groups of each SNP.

### Human Clincial Sample

There are 315 healthy Caucasian volunteers in the current study, 141 male and 174 female, and the mean age 32.7±13.0 years (age range 18-86 years). Results examined using passed clinical and imaging quality control measures. Exclusion criteria included a current or past history of head trauma or neurological/psychiatric disorders, previous medical treatment or medication at the time of participation pertaining to metabolism or blood flow, or history of drug/alcohol abuse according to DSM-IV criteria (following a Structured Clinical Interview, SCID-IV, (1)). All participants underwent functional magnetic resonance imaging (fMRI) while performing an incidental encoding and memory retrieval task. All participants gave written informed consent for both sessions, approved by the National Institute of Mental Health Institutional Review Board. The details of subject recruitment, screening, medical review and scanning and cognitive task administration and scoring are described in detail elsewhere (Rasetti et al^3^).

### Human Clinical Neuropsychological Testing

All clinical subjects underwent a battery of neuropsychological testing to assess their cognitive function before the fMRI study. Following previous publication (Museet al^4^), we investigated Logical Memory (LM) I and II with current samples for WWC1, PRKCG, and PRKCZ polymorphisms. Fisher’s r-to-z transform in the R package psych was applied on neuropsychological test data before conducting comparison tests.

### Human Clinical fMRI Paradigm

Blood oxygen level-dependent (BOLD) fMRI was collected during the encoding and subsequent retrieval of semantically unrelated aversive and neutral complex scenes selected from the International Affective Picture System (Lang et al^5^). For both the encoding and retrieval sessions, the scenes were presented in a blocked fashion with two blocks of aversive or neutral scenes alternating with resting-state blocks. The order of presentation of the aversive vs. neutral scene blocks was counterbalanced across subjects. During each experimental block, six scenes of similar valence (neutral or aversive) were presented serially to participants for three seconds each. Each resting block spanned 18 seconds, during which subjects were instructed to attend to a fixation cross located in the middle of the screen. Before starting each session (encoding and retrieval), subjects were given verbal instructions and asked to verbally indicate to the task administrator that they understood the instructions. Additionally, the task instructions were presented on the screen for two seconds at each session’s beginning. For encoding, participants were instructed to determine whether a presented picture depicted an indoor scene or outdoor scene by button presses (right button for “Outdoor scene” or left button for “Indoor scene”). For retrieval, participants were presented with 50 percent of the scenes from the encoding session (old) and 50 percent novel scenes (new). They were instructed to determine whether or not they had previously seen each picture in the indoor/outdoor task by button presses (right button for “Yes” or left button for “No”) using their dominant hand. Subjects were neither encouraged nor discouraged from guessing for either session, and only subjects with performance above chance (> 50 % accuracy) during both sessions were included in the final analysis. Each session’s total scan time was 5 minutes and 40 seconds, which included the stimulus blocks (4 aversive and 4 neutral), resting blocks (9), and instruction screen. Task performance (% correct responses) and response time (seconds) were recorded for each session.

### Human Clinical fMRI Data Acquisition

BOLD fMRI was performed using a GE Signa 3-Tesla Signa scanner (Milwaukee, WI) with a gradient EPI sequence (24 axial slices; gap = 1 mm; thickness = 4 mm; TR/TE = 2000/28; scan repetitions = 170; flip angle = 90°; field of view = 24 cm; matrix = 64 x 64). Scanning parameters were selected to optimize the BOLD signal’s quality while maintaining a sufficient number of slices to acquire whole-brain data.

### Human Clinical Neuroimaging Data Analysis

#### Preprocessing

Images were processed using SPM12 software (www.fil.ion.ucl.acuk/spm). Image preprocessing was performed as follows: image realignment to the first scan of the run to correct for head motion, spatial normalization to the Montreal Neurological Institute standard template using an affine model, and smoothing using an isotropic Gaussian filter (8 mm^3^ full-width, half-maximum kernel). Each dataset was carefully screened for data quality by performing a visual inspection for image artifacts, an estimation of indices for ghosting artifacts, a signal-to-noise ratio (SNR) calculation across the time series, and a signal variance statistic across individual sessions. Data from participants with head motion greater than 3 mm, head rotation greater than 2°, imaging artifacts, and/or low SNR were excluded.

#### 1st Level Processing

For the functional neuroimaging data, fMRI responses were modeled using the general linear model with a canonical hemodynamic response function convolved to a boxcar function for the length of the block, normalized to the global signal across the whole brain, and temporally filtered to remove low-frequency signals (<84 Hz). Condition effects at each voxel for each subject were created using a *t*-statistic in SPM12, producing statistical images of significantly-activated brain regions for the following contrasts of interest: neutral encoding > baseline and neutral retrieval > baseline.

#### Group Level Analysis

Individual subject first-level contrast images related to the encoding of neutral scenes were entered into second-level full factorial model (ANCOVA) to evaluate the effect of WWC1, PRKCG, and PRKCZ polymorphisms on brain activation changes along age, as well as the interactions of these polymorphisms in the brain regions that were activated in fMRI task. Since our primary goal was to explore the potential way in which age-related changes in neuronal activity are affected by the WWC1, PRKCG, and PRKCZ genes, and their interactions, current analyses were limited to neural activation within the hippocampal formation (HF). A anatomical mask including bi-lateral hippocampal and parahippocampal regions was created using the WFU Pickatlas toolbox (version 1.04; Functional MRI Laboratory at the Wake Forest University School of Medicine, Winston-Salem, North Carolina; http://fmri.wfubmc.edu/). We conducted small volume correction within HF, and clusters including at least one volxel passing FDR<0.1 were reported.

#### Extracted ROI Analysis

For each significant cluster, a sphere with 6mm radius around the peak and withith HF mask was defined as the region of interest (ROI). The mean value from each individual’s first level contrast image was extracted from ROI for further analysis. Main effect and interaction effect of polymorphisms were evaluated in R with general linear model using the extracted values. IQ were used as covariates since IQ shows signifincat association with KIBRA SNPs rs17070145.

##### Statistical Test Methods

The effect of SNPxAge interaction was evaluated using likelihood ratio test (lrtest) in lmtest package in R as shown in the following formula:

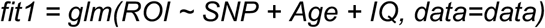

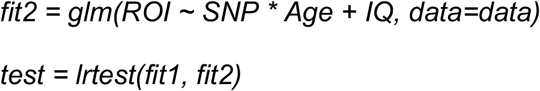

The comparison of interaction of SNPxAge between genotype groups was evaluated using lstrends in lsmeans package in R as shown in the following formula:

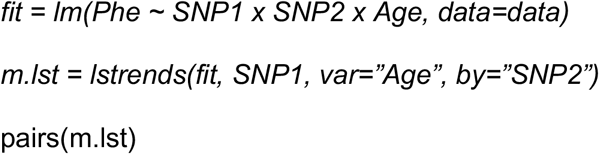

Based on the estimate and SE reported from the pairs function, we did Welch’s t test to compare interaction effects between SNP1 and age within two genotype groups of SNP2.

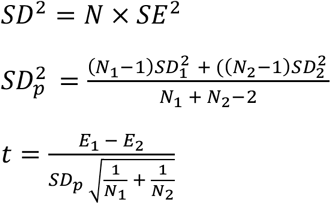

P values were converted using pt function in R.

## Acknowledgements

We thank Drs. Meaghan Morris, Graham Diering, and Victor Anggono for critical reading of the manuscript, and thank all the members of R.L.H. laboratory for discussion and support. This work was supported by NIH R01NS036715 to R. L. H., and the Johns Hopkins University Science of Learning Institute postdoc fellowship to M.T.

## Author Contributions

M.T., D.W. and R.L.H. designed experiments. M.T., A.G., H.G., and R.J. performed biochemistry analysis. D.C. and D.W. analyzed human subject data. M.T. and R.L.H. wrote the manuscript with input from all authors.

## Additional Information (Containing Supplementary Information Line (If Any) And Corresponding Author Line)

## Data availability

All requests for raw and analyzed data and materials are promptly reviewed by the Johns Hopkins University School of Medicine or the Lieber Institute of Brain Development to verify if the request is subject to any intellectual property or confidentiality obligations. Human-related data not included in the paper might be subject to confidentiality. Any data and materials that can be shared will be released via a Material Transfer Agreement. Raw data used in the generation of Figs. 1–4 and Supplementary Fig. 1 are available upon request.

## Extended Data Figure/Table Legends

**Suppl. Table 1.** Statistical results of NeuroPsych analysis.

| Gene | Test |  | N | A_Imm | A_Delay |
| --- | --- | --- | --- | --- | --- |
| KIBRA | rs17070145 |  | CC/T: 116/139 | 0.2138 | **0.0159 \*** |
|  | rs17070145 x Age |  |  | 0.1585 | 0.0565 |
| PRKCG | rs12151321 |  | A/CC: 220/30 | 0.381 | 0.3867 |
|  | rs12151321 x Age |  |  | 0.7494 | 0.5902 |
|  | rs12151321=A | rs17070145 | CC/T: 103/117 | 0.1422 | **0.0190 \*** |
|  |  | rs17070145 x Age |  | 0.1451 | **0.0271 \*** |
|  | rs12151321=CC | rs17070145 | CC/T: 11/19 | 0.4440 | 0.8050 |
|  |  | rs17070145 x Age |  | 0.6572 | 0.9492 |
| PRKCZ | rs2803349 |  | C/TT: 187/61 | 0.5546 | 0.9339 |
|  | rs2803349 x Age |  |  | 0.5192 | 0.3240 |
|  | rs2803349=C | rs17070145 | CC/T: 83/104 | 0.0929 | **0.0026 \*\*** |
|  |  | rs17070145 x Age |  | 0.2776 | 0.1017 |
|  | rs2803349=TT | rs17070145 | CC/T: 27/34 | 0.9770 | 0.9190 |
|  |  | rs17070145 x Age |  | 0.5791 | 0.7810 |

\*: p < 0.05
\*\* : p < 0.01

**Supplementary Table 2.**
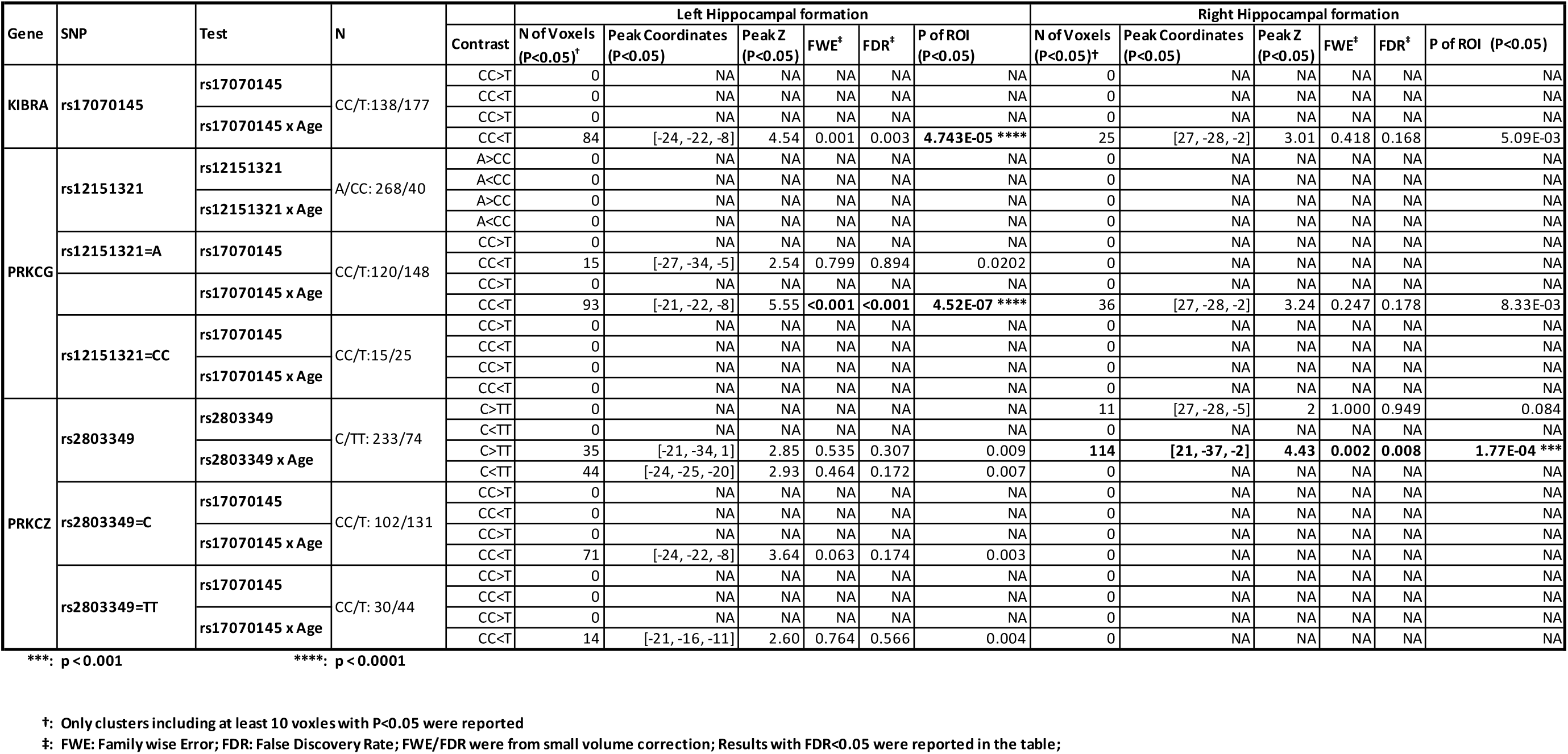
Statistical analysis of MTL activities.

## Supplementary Figures and Legends

**Supplementary Fig 1.**
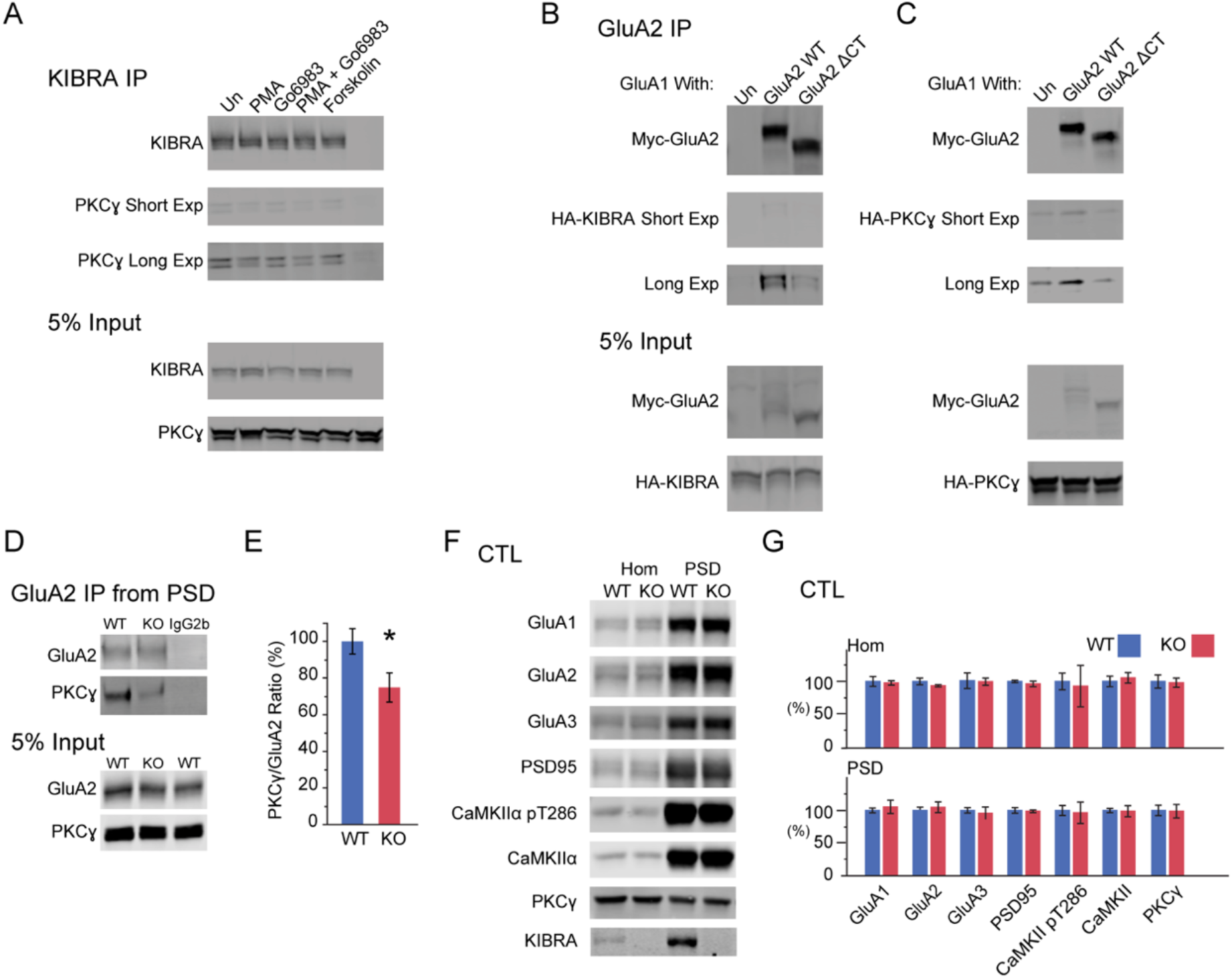
**A**. Representative western blots showing co-immunoprecipitation of PKCγ with KIBRA when co-expressed in HEK293T cells. Eighteen hours after transfection, HEK293T cells were treated with the chemical labeled for 5 minutes and lysed in lysis buffer containing 1% NP40 and 0.5% Deoxycholate Sodium. PMA 2 μM, Go6983 10 μM, Forskolin 50 μM. **B**. Representative western blots showing co-immunoprecipitation of PKCγ with GluA2 decreased when GluA2 C-terminus tail (GluA2 ΔCT) is truncated. PKCγ co-expressed with GluA2 WT or GluA2 ΔCT in the presence of WT GluA1 in HEK293T cells for 18 hours. **C**. Representative western blots showing co-immunoprecipitation of KIBRA with GluA2 decreased when GluA2 C-terminus tail (GluA2 ΔCT) is truncated. KIBRA co-expressed with GluA2 WT or GluA2 ΔCT in the presence of WT GluA1 in HEK293T cells for 18 hours. **D**. Representative western blots showing co-immunoprecipitation of PKCγ with GluA2 from PSD fraction of mouse forebrain. **E**. Quantification of **D**. n = 6. **F**. Representative western blots of proteins from Hom and PSD isolated from gender-mated WT and KIBRA KO littermate hippocampus in home cage. **G** Quantification of **F**. WT: n = 7; KO: n = 11. \**p* < 0.05.

**Supplement Fig 2.**
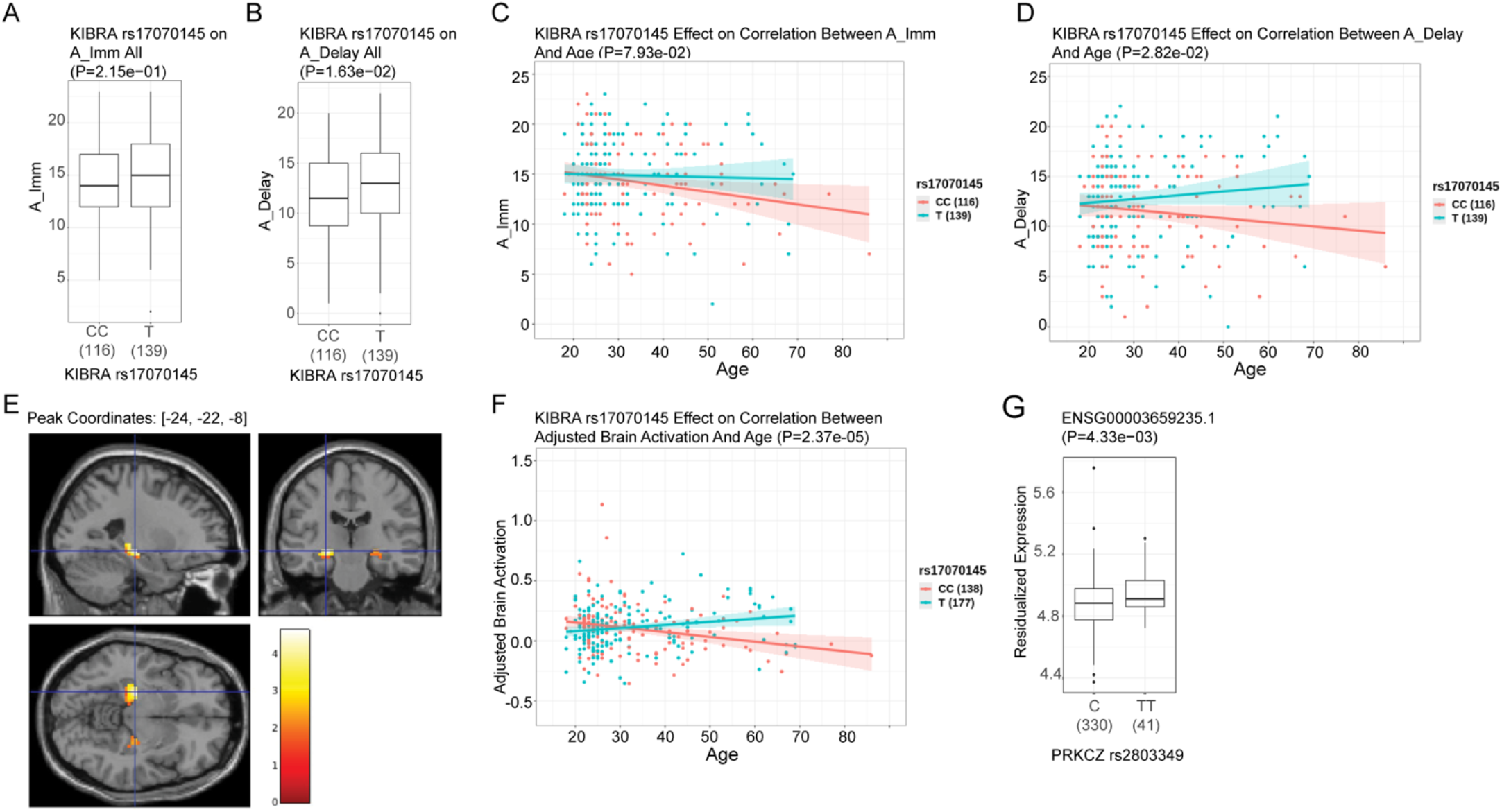
The allelic association of the *KIBRA* gene with human memory performance. **A-D:** Effect of KIBRA SNP rs17070145 on memory performance **A:** no effect of *KIBRA* SNP rs17070145 on immediate recall **B**. Significant effect of *KIBRA* SNP rs17070145 on 30-minute delayed recall (*p* = 0.0163), with T carriers performing better than individuals with homozygous C allele. **E**, **F**. Association of age with brain activation under episodic memory task affected by KIBRA SNP rs17070145 **G.** eQTL effect on expression of *PRKCZ*.

